# Differential innate immune response of endometrial cells to porcine reproductive and respiratory syndrome virus type 1 versus type 2

**DOI:** 10.1101/2022.10.10.511536

**Authors:** Muttarin Lothong, Dran Rukarcheep, Suphot Wattanaphansak, Sumpun Thammacharoen, Chatsri Deachapunya, Sutthasinee Poonyachoti

## Abstract

Modification of cellular and immunological events due to porcine reproductive and respiratory syndrome virus (PRRSV) infection is associated with pathogenesis in lungs. PRRSV also causes female reproductive dysfunction and persistent infection which can spread to fetus, stillbirth, and offspring. In this study, alterations in cellular and innate immune responses to PRRSV type 1 or type 2 infection, including expression of PRRSV mediators, mRNA expression of toll-like receptor (TLRs) and cytokine, and cytokine secretion, were examined in primary porcine glandular endometrial cells (PGE). Cell infectivity as observed by cytopathic effect (CPE), PRRSV nucleocapsid proteins, and viral nucleic acids was early detected at two days post-infection (2 dpi) and persisted to 6 dpi. Higher percentage of CPE and PRRSV positive cells were detected in type 2 infection. PRRSV mediator proteins, CD151, CD163, sialoadhesin (Sn), integrin and vimentin, were upregulated following type 1 and type 2 infection. *CD151*, *CD163* and *Sn* were upregulated by type 2. Both PRRSV types upregulated *TLR1* and *TLR6.* Only type 2 infection upregulated *TLR3,* but downregulated *TLR4* and *TLR8*. By contrast, both types upregulated TLR4 and downregulated TLR6 protein expression. *Interleukin* (*IL*)-*1β*, *IL*-*6* and *tumor necrotic factor* (*TNF*)-*α* were upregulated by type 2, but *IL*-*8* was upregulated by type 1. Both PRRSV type 1 and 2 stimulated IL-6 but suppressed TNF-α secretion. In addition, IL-1β secretion was suppressed by type 2. These findings reveal one of the important mechanisms underlying the strategy of PRRSV on innate immune evasion in endometrium which is associated with the viral persistence.

**Author Summary:** Widely prevalence of porcine reproductive and respiratory syndrome virus (PRRSV) remains the leading cause of huge economic losses to the global swine industry. Due to an infection of macrophages, PRRSV can persist in animals for extended periods of time associated with long-lasting reproductive disorders. Modification of cellular and immunological responses to PRRSV infection which may be related with the pathogenesis of reproductive disorders remains unclear. Herein, direct PRRSV infection of primary porcine glandular endometrial epithelial cell culture (PGE) demonstrated that PRRSV type 1 and 2 upregulated the protein expression of PRRSV mediators correlated with cell persistence of PRRSV. However, TLR and cytokines gene expression, and cytokine secretion were differentially modulated in response to PRRSV type 1 vs. type 2. Our study provides new insights into the cellular mechanism associating with PRRSV persistence in the endometrial cells and the underlying interaction of virus with the host.

## Introduction

Porcine reproductive and respiratory syndrome virus (PRRSV) disease is one of the most economic issue that affects the swine industry [1]. Massive reproductive disorders in sows causing perinatal losses and respiratory distress in piglets is the main impact of PRRSV outbreak [2]. Increased expenses associated with recirculation of PRRSV infection, i.e., confirming PRRSV status and treatment of secondary infections have an indirect impact on the cost of production [1].

PRRSV is a positive single-stranded RNA virus belonging to the *Arteriviridae* family. Two distinct strains, PRRSV type 1 and type 2, are identified and share approximately 60% genetic identity [3]. The virulence, clinical severity, antigenic and immunological diversity are relevant to genetic variability of PRRSV strains [3, 4].

In reproductive organs, PRRSV can be transmitted via transplacental viral shedding leading to aborted fetus and/or weak born piglet [5]. It has been suggested that PRRSV replication can occur at the implantation site [6], although PRRSV is highly restricted to porcine alveolar macrophages (PAMs) [7, 8].

PRRSV mediators are the major determinant of cellular susceptibility. At least five molecules have been described as PRRSV mediators, such as CD163, Sn, CD151, integrin and vimentin [9, 10]. These receptors play significant roles in PRRSV infection, including virus binding, internalization, uncoating and replication [11]. Previous study in primary porcine glandular endometrial cells (PGE) demonstrates a low expression level of PRRSV receptors, CD163 and Sn; however, PRRSV can induce CPE and damage PGE [12]. Moreover, high existence and release of PRRSV from PGE was gradually increased with time suggesting the susceptibility or recirculation of PRRSV in the infected PGE [12]. Overexpression or modification of PRRSV receptor cDNA in PRRSV non-permissive cell lines can generate the infectious progeny virus from those cells [13]. This raised the possibility that PGE may express alternative PRRSV receptors, in which PRRSV can infect and process host cellular response. The modification of PRRSV receptor expression associated with the increased PRRSV susceptibility and replication remains to be investigated.

Innate immune response is primarily responsible for an immediate protection against pathogen invasion and subsequently activates the adaptive immune response. The innate immune response is mediated through host pattern recognition receptors (PRRs), i.e., toll-like receptors (TLRs). The TLRs 1-9 are expressed in immune cells and non-immune cells like endometrial epithelial cells [14]. Endosomal TLRs, TLR3, TLR7, TLR8, and TLR9 detect internalized viral nucleic acid and promote the production of anti-viral cytokines such as type I interferons (IFN), IFN-α and IFN-β [15]. Membranous TLRs, TLR2 and TLR4 generally recognize bacterial structural proteins but not the invading viral particles. TLR and cell signaling system mediate host-viral PRRSV interaction by inducing the production of pro-inflammatory cytokines, such as IL-6, IL-8 and TNF-α in MARC-145 and PAMs [16]. PRRSV infections are characterized by prolonged viremia and complication from immunosuppressive effects because of the upregulation of *IL*-*10* and *IL*-*1β* expression and reduction of *IFN*-γ in porcine polymorphonuclear cells [17, 18]. Thus, the cellular modification of host immune system is relevant to both pathogenesis and host protection of PRRSV. The effect of PRRSV infection on the local innate immunological responses of the reproductive system is associated with reproductive failure is still in question.

In the present study, we employed the PGE infected with PRRSV from our previous study [12] to determine an *in vitro* cellular response in association with cytopathic effect (CPE) and viral replication pertaining to the expression of aforementioned PRRSV mediators, TLRs and pro-inflammatory cytokines secretion following PRRSV type 1 or type 2 infection.

## Results

### PRRSV type 2 induced CPE and viral existent in PGE

As shown in Fig. 1, PGE infected with PRRSV type 1 or type 2 showed CPE, including syncytial formation (Sc; Fig. 1A) and vacuolization (Vc; Fig. 1A) as early as 2 dpi. The CPE area at 2 dpi were significantly extended to 20-30% in both type 1 and type 2 inoculated PGE as compared to mock and uninfected PGE (Fig. 1B). A greater extent of CPE area (40%) was found in type 2 inoculated PGE at 4 dpi, but it was recovered at 6 dpi (Fig. 1B). By contrast, the CPE area produced by type 1 were not significantly different from that by the mock and uninfected groups at 6 dpi (Fig. 1B). Corresponding to CPE formation, immunoreactivity of PRRSV envelop protein (GP5; Fig. 1C) was early observed at 2 dpi and remained up to 6 dpi. Type 2 inoculation increased the PGE immunoreactive cells about 3-4 times higher than type 1 inoculation (Fig 1C, *p* < 0.05).

**Figure 1.**
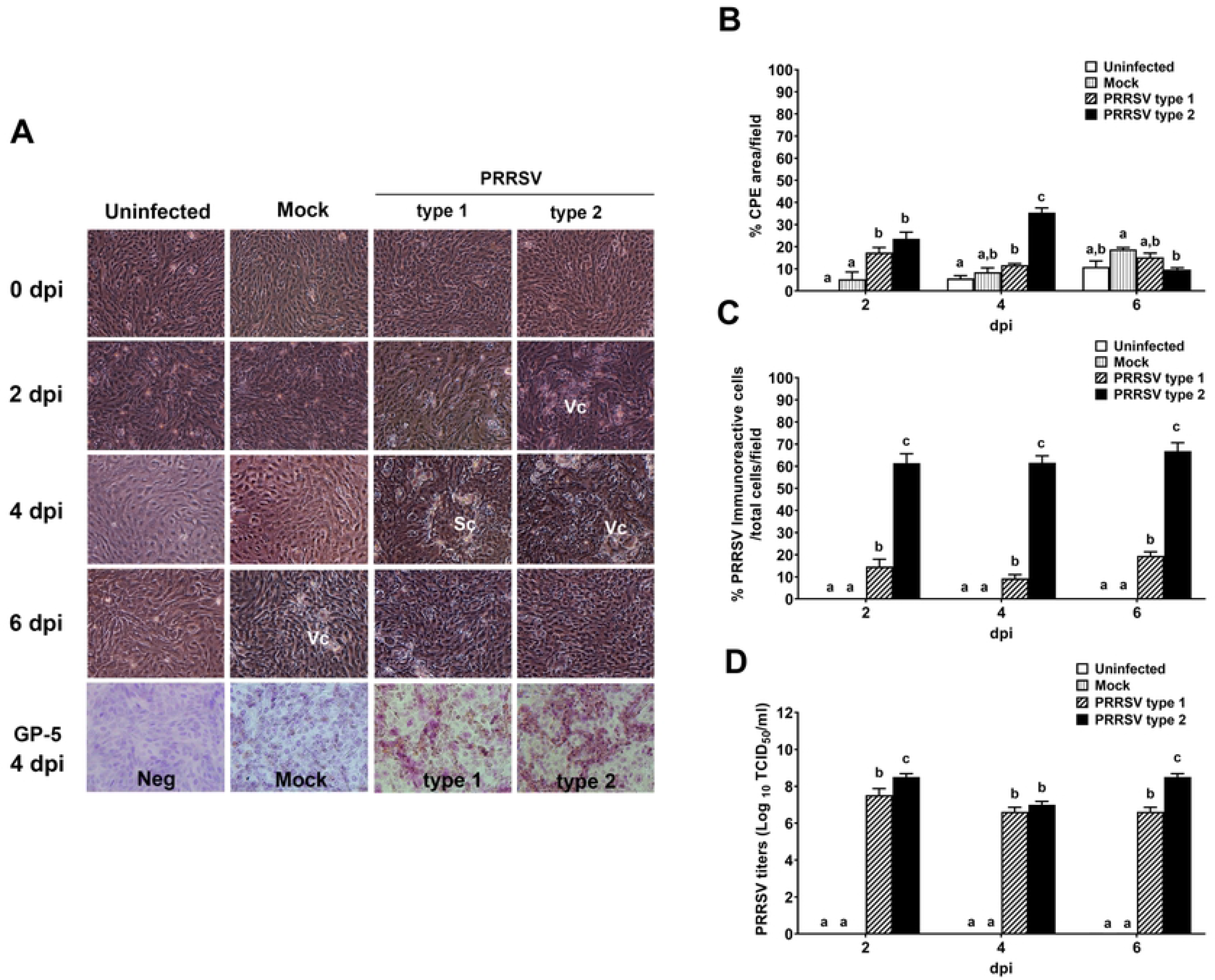

The PRRSV existing in type 2-inoculated PGE was 10^9^ TCID_50_/mL which were a ten-fold higher than type 1 at 2 and 6 dpi (Fig. 1D; *p* < 0.05). However, the PRRSV titers were equivalent in both type 1 and type 2 infected PGE at 4 dpi (Fig. 1D; *p* > 0.05).

### PRRSV type 2 upregulated *CD151*, *CD163* and *Sn* mRNA expression

The mRNA expression of PRRSV mediators, *CD151*, *CD163*, *Sn, integrin,* and *vimentin,* was determined at 4 dpi to examine whether they might be a target involved in PRRSV infection and existent. The normal uninfected PGE expressed a relatively higher level of *vimentin* (3-fold) and low level of *CD151*, *CD163*, *Sn* and *integrin* (0.5-fold) as normalized to *GAPDH* (Fig 2A). At 4 dpi, only type 2 infection was found to upregulate the expression of *CD151, CD163* and *Sn* by 4-60-fold (*p* < 0.05, Fig 2B) with no effects on *integrin* or *vimentin*. However, both mock and type 1 infection could not produce any effects on the mRNA expression of PRRSV mediators tested (*p* > 0.05; Fig 2B).

**Figure 2.**
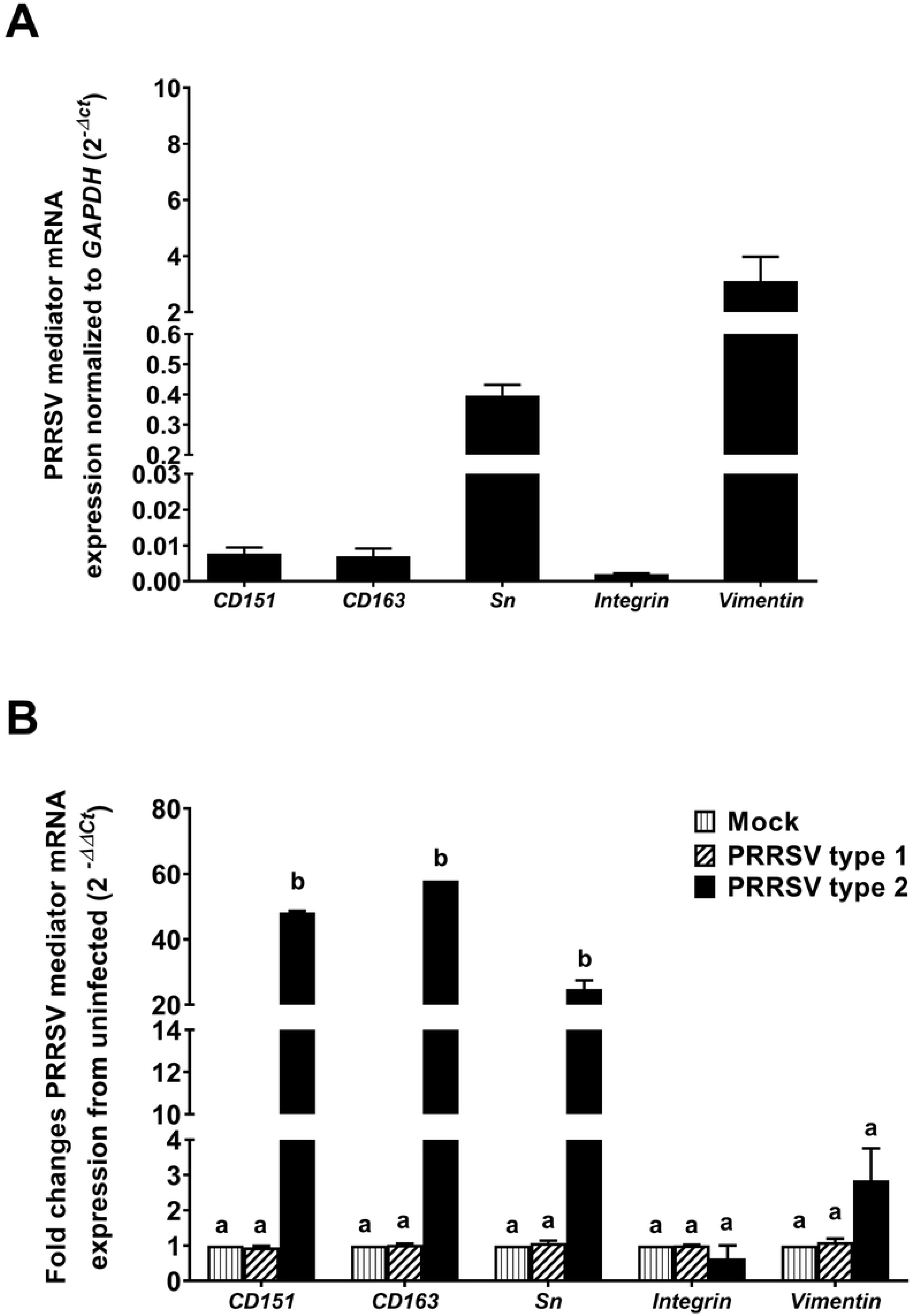

### PRRSV type 1 and type 2 upregulated cellular expression of PRRSV mediators

Apart from mRNA expression, cellular localization of PRRSV mediator proteins at 4 dpi was further evaluated using immunohistochemistry in PGE. In uninfected group, at 0 dpi, the immunoreactivity of CD151, CD163, Sn and integrin was rarely detected, whereas that of vimentin was demonstrated about 10% (Fig. 3A). The immunoreactive CD151, CD163, Sn or integrin was distributed in the cytoplasm of mock or PRRSV infected PGE (Fig. 3A). In contrast, the vimentin immunoreactivity which had fiber-like characteristics was observed in the uninfected and mock PGE. Analysis of expression area of PRRSV mediators was not difference between the uninfected and mock PGE (*p* > 0.05; Fig 3A and 3B). Both type 1 and type 2 markedly upregulated the expression of all PRRSV mediators. In this result, the increases in CD151, CD163, Sn, and integrin expression induced by type 2 was higher than type 1 (*p* < 0.05; Fig 3B).

**Figure 3.**
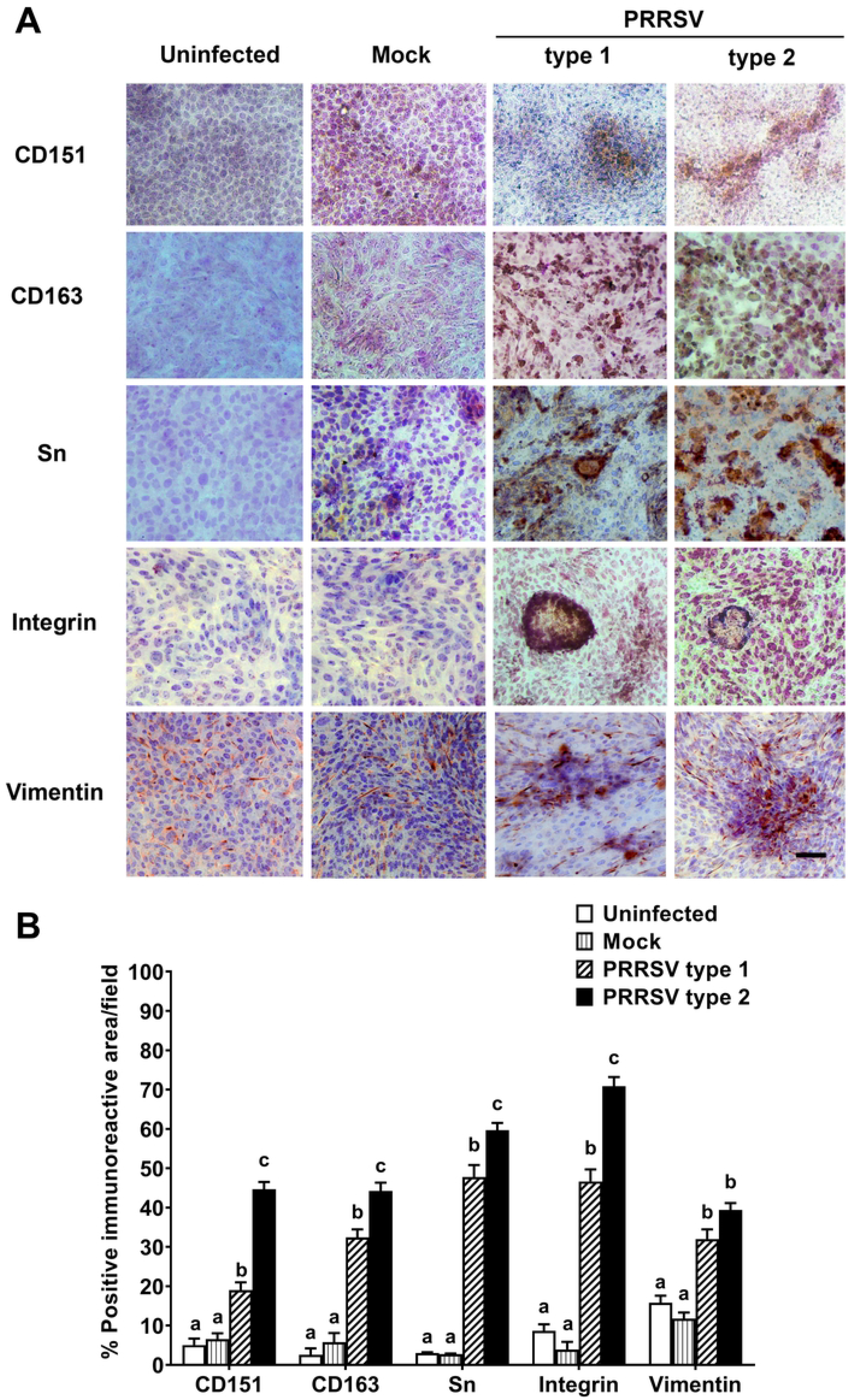

### PRRSV type 1 and type 2 upregulated TLR6 and downregulated TLR4 expression

Changes in mRNA and protein expressions of TLRs 1-10 by PGE were also determined following 4 dpi. RT-PCR results of *TLR1*-*10* gene expression in Fig 4A showed that PRSSV type 1 and type 2 infection upregulated *TLR1* and *TLR6*, while only type 1 upregulated *TLR3* expression (*p* < 0.05; Fig 4A). In contrast, *TLR4* and *TLR8* was downregulated in type 2 infected group (*p* < 0.05; Fig 4A). Most consistent with mRNA expression, the protein expression of TLR1, TLR3, TLR4, TLR6 and TLR10 in uninfected and mock infected groups was not different (*p* > 0.05, Fig 4B and 4C). Both PRRSV type 1 and type 2 upregulated TLR6 by 2-fold from uninfected group whereas they suppressed TLR4 expression (*p* < 0.05; Fig 4B).

**Figure 4.**
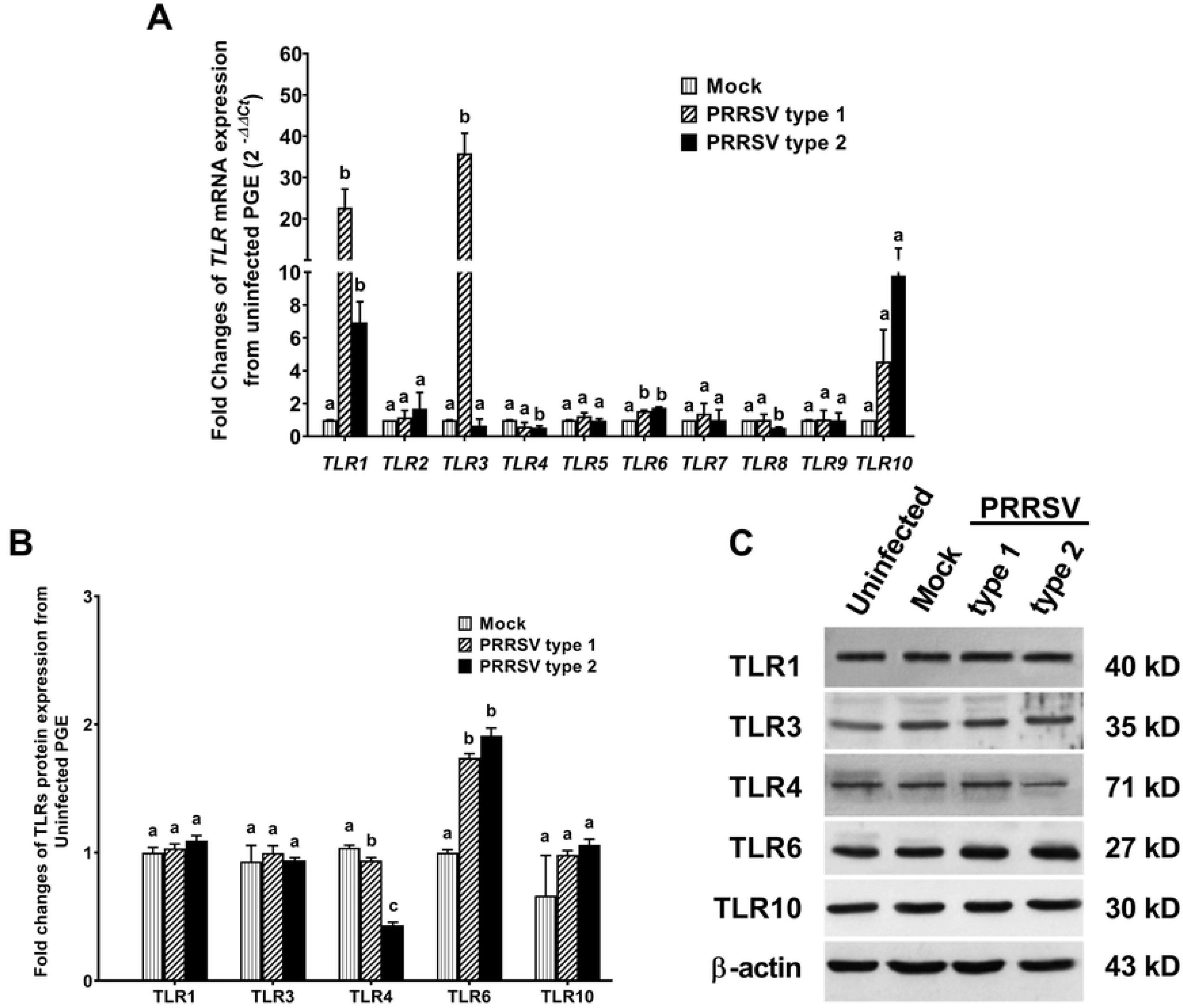

### PRRSV type 1 and type 2 upregulated IL-6 and downregulated TNF-α

The mRNA expression of cytokines at 4 dpi and accumulated cytokine secretion at 6 dpi were determined following PRRSV inoculation in PGE. The mRNA expression of cytokines was not different between uninfected and mock infected groups (*p* > 0.05, Fig 5A). *IL-8* was upregulated by type 1, and *IL-1β, IL-6* and *TNF-α* were upregulated by type 2 infection (*p* < 0.05; Fig 5A).

**Figure 5.**
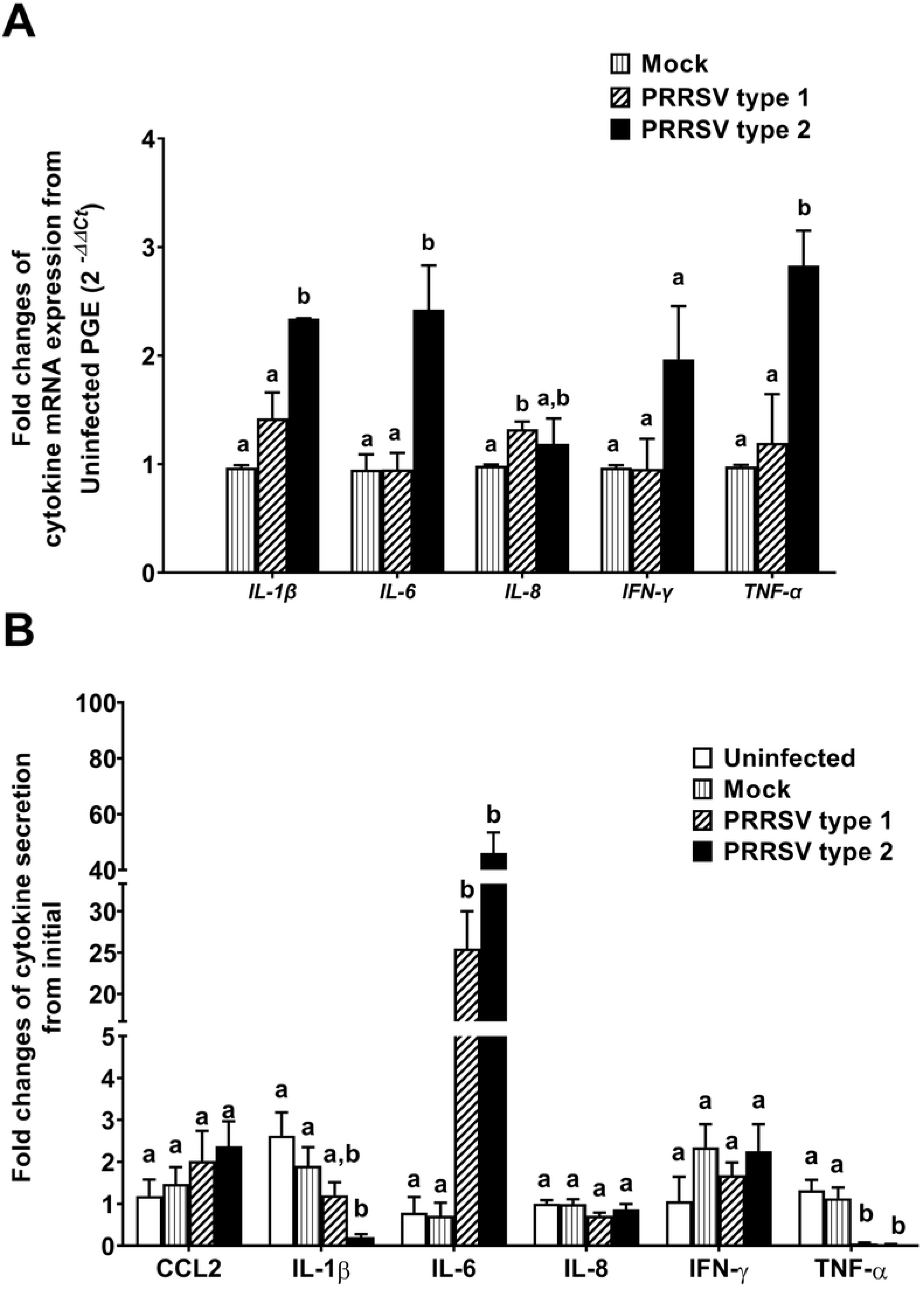

Both PRRSV type 1 and 2 stimulated IL-6 but suppressed TNF-α secretion (*p* < 0.05; Fig 5B). The accumulated secretion of IL-1β was additionally suppressed after infection with type 2 (*p* < 0.05; Fig 5B). CCL2, IL-8 and IFN-γ secretions were not different among groups (*p* > 0.05; Fig 5B). Moreover, there was no difference in cytokine secretion between mock and uninfected groups (*p* > 0.05; Fig 5B).

## Discussion

Besides alveolar macrophages, the natural cell trophism of PRRSV, PGE has been recently reported as an alternative target of PRRSV, which was preferentially infected via the apical side of PGE monolayer [12]. In relation to the previous study, PGE infectivity as observed by increases in CPE, PRRSV mediators and viral titers were found to be associated with changes in cellular innate immune system, toll-like receptor expression, and cytokine expression and secretion following the apical PRRSV infection in the present study. Both PRRSV type 1 and type 2 infection not only produced the cytotoxic effects but also persisted in PGE up to 6 dpi. However, different cellular responses of PGE to PRRSV type 1 and type 2 infection were observed.

In the current study, PRRSV type 2 infected PGE displayed more CPE and higher percentage of PRRSV positive cells than type 1. Thus, more cellular destruction and PRRSV existence caused by type 2 infection indicated the higher virulence to PGE than that by type 1. This finding was consistent with natural infection, which PRRSV type 2 revealed more severity of respiratory distress than type 1 [3]. However, the natural infection or *in vivo* intranasal inoculation with type 1 or type 2 could not demonstrate the different virulence in reproductive signs, i.e., viral load in fetus or maternal-fetal interface, the number of embryonic death or PRRSV-positive litters [19]. In our previous studies, apical route of PRRSV infection alters the permeability and viability of PGE [12, 20]. However, the basolateral route of PRRSV infection at endometrium occurred following viremia also causes reproductive disorders [21]. Therefore, the different route of PRRSV infection which appears to produce severity in reproductive tissue dysfunction should be taken into consideration and confirmed by further study.

Our study is the first report to indicate the upregulated cellular expression of PRRSV mediators in response to PRRSV infection in porcine endometrial cells. PGE infected with PRRSV type 2 induced higher degree of upregulated mRNA expressions of *CD151*, *CD163* and *Sn* which were associated with the high titers of PRRSV persistence and release from infected cells. The increased PRRSV mediators in PGE may play a substantial role in the sensitivity and persistence to the consecutive PRRSV infection.

Besides, CD151 expression was found to be significant higher in endometrioid carcinoma, sarcoma or carcinosarcoma [22]. Interaction between CD151 and integrin promoted outgrowth in human embryonic carcinoma cell line NT2N [23]. Therefore, upregulated CD151 may explain the proliferative effects of PRRSV type 2 infection in PGE [20]. Moreover, the infected PGE was associated with severe membrane integrity dysfunction [20]. Thus, modulation of PRRSV mediators following PRRSV infection may be the underlying pathogenesis of endometrial function leading to the reproductive failure in sows.

TLRs and cytokines are the key components of mucosal innate immune responses in female reproductive tract. Upregulated TLR3, TLR7, and TLR8 expressions were correlated to PRRSV virulence and clinical signs in highly pathogenic PRRSV-infected pigs [24]. Differential regulation on *TLR1-10* expression was also observed following type 1 or type PRRSV infection in PGE. Type 1 infection produced stimulatory effects on *TLRs*, *TLR1, TLR3* and *TLR6* which seems not to be correlated with virulent characteristics found in the present study. Although the upregulated *TLR1* and *TLR6* are not described as virulent characteristics [24], our findings that PRRSV type 2 upregulation of *TLR1* and *TLR6* and downregulation of *TLR4* and *TLR8* are likely to produce high infectivity, virulence, and persistence in the reproductive epithelial cells PGE. Among the target genes of PRRSV infected PGE including *TLR1, TLR3*, *TLR4*, *TLR6* and *TLR8* were indicated, only TLR4 and TLR6 protein expression were changed by PRRSV infection.

Upregulated TLR protein expression in PRRSV infected pigs contributes to disease progression or the severity of clinical respiratory signs because of an excessive response of pro-inflammatory cytokine via PAMPs/TLR signaling system [25]. Constitutive secretion of IL1β, IL-6, IL-8 and TNF-α which is mediated through TLRs activation by TLRs ligand, including poly I:C dsRNA simulating viral nucleic acids has been previously demonstrated in PGE [26]. Secretion of CCL2 and IFN-γ by PGE was additionally demonstrated in this study; however, secretion of IL-10 and IFN-α was absence in all groups (data not shown). Thus, the presence of these innate immunity-related molecules indicates the ability of PGE to interact with the viral pathogens and establish a major part of the innate immune response of endometrial cells.

In PGE, PRRSV type 2, but not type 1 infection, induced *IL-1β, IL-6, IL-8* and *TNF-α* mRNA expression at 4 dpi, which revealed the highest CPE area of infected PGE. The recent findings agree with the virulent effects of PRRSV type 2 in *in vitro* studies of PAMs and peripheral blood mononuclear cell (PBMCs) [18, 25, 27].

In general, pro-inflammatory cytokines IL-1β and TNF-γ are released by host to encounter invasive pathogens including viruses. The downregulated TNF-α synthesis and secretion by PRRSV infection in PGE were consistent to the study of PAMs infected with PRRSV type 2 [28]. Incubation with recombinant TNF-α was reported to reduce existent of PRRSV [28]. In PRRSV-infected PGE, particularly type 2, the increased viral existence at 2-6 dpi (Fig. 1D) was concurrent to the decreased IL-1β and TNF-α.

The suppressed TNF-α induced by PRRSV has been indicated to carry out by Nsp1 by inhibiting the activation of NF-κB and Sp1 transcription factors on the promoter region of *TNF-γ* [29], whereas upregulated IL-β production in PAMs induced by PRRSV was mediated via TLR4/MyD88 pathway [30]. Therefore, the reduced IL-1β secretion in PGE by type 2 may be due to the suppressive effect of PRRSV on TLR expression. This was evidenced by the highest degree of TLR4 suppression and IL-1β inhibition in type 2 infected PGE. As the expression level of IL-1β was related to PRRSV clearance [31], the reduced TLR4/IL-1β system and TNF-α production by PRRSV might be the strategy of virus to escape the obliteration of host cells. Herein, the decreased CPE with persistence of PRRSV type 2 at 6 dpi may support these assumptions. Taken together, the downregulated TNF-α and IL-1β secretion by PRRSV in immune and PGE cells reflects a poor innate immune response which leads to secondary infection by other microbial pathogens [32].

Additionally, the largely increased IL-6 secretion was found in response to PRRSV infection. Excessive expression of IL-6 followed by the increased expressions of pro-inflammatory cytokines and chemokines, including IL-1β, IL-12, IL-8, and TNF-α was relevant to the severe lung injury and damages of lymphoid organs in highly pathogenic PRRSV (HP-PRRSV) [33]. It is likely that type 2 induced higher damages of PGE in our previous [20] and current studies may be explained by the increase in *IL-1β*, *IL-6* and *TNF-α* mRNA expression.

Apart from immune function, pro-inflammatory cytokines play roles in pregnancy and parturition. The locally increased IL-1β and TNF-α from neutrophil infiltrated uterine are required for muscle contraction and cervical ripening during parturition [34]. However, application of IL-1β and/or TNF-α caused the dissolution of collagen fibers, stromal edema, and severe inflammation in the cervix of guinea pigs [35]. Exposure or infusion of IL-1β and TNF-α caused the defects associated with peripartum intrauterine inflammation, abnormal lung development associated with bronchopulmonary dysplasia, and brain injury [34]. Thus, the modulation of cytokine synthesis and release by PRRSV infection in PGE affecting the reproductive infertility needs a further study.

## Materials and Methods

### Reagents and materials

Cell culture-grade reagents of Ringer’s solution, ethanol, isopropanol, H2O2 and methanol were purchased from Sigma Chemical Co., (St Louis, MO, USA). Dulbecco’s Modified Eagle’s Medium (DMEM), Dulbecco’s PBS, fetal bovine serum (FBS), collagenase type I, 0.05% trypsin-0.53 mM ethylenediaminetetraacetic acid (EDTA), kanamycin, penicillin-streptomycin and fungizone were purchased from Gibco BRL (Grand Island, NY, USA).

### Antibodies

Rabbit polyclonal antibody (pAb) against PRRSV envelop glycoprotein GP5 (Biorbyt Ltd., Cambridge, UK; dilution 1:100) was used to detect PRRSV protein. To localize PRRSV mediators, antibodies with the final dilution including rabbit pAb anti-human CD151 (Abcam, Waltham, MA, USA; 1:250), goat pAb-anti-porcine CD163 (Santa Cruz biotechnology, Dallas, TX, USA; 1:25), mouse monoclonal Ab (mAb)-anti-sialoadhesin (Bio-rad, Inc., Hercules, CA, USA; 1:250), goat pAb anti-integrin-α3 (Santa Cruz biotechnology, Dallas, TX, USA; 1:25) and mAb anti-vimentin (Santa Cruz biotechnology, Dallas, TX, USA; 1:250) were used for immunostaining. To assess the expression of toll-like receptors, mAb anti-TLR1, mAb anti-TLR3, mAb anti-TLR4, mAb anti-TLR6, and mAb anti-TLR10 were purchased from Santa Cruz biotechnology (Dallas, TX, USA). Other antibodies including mAb anti-β-actin, HRP-conjugated anti-mouse IgG, and mAb anti-goat IgG were purchased from Bio-rad (Hercules, CA, USA).

### PGE cell culture

PGE were isolated from Thai crossbred commercial 4–6 months old gilts provided by a government-qualified slaughterhouse in Bangkok, Thailand, following a previous protocol [36]. Briefly, the uterine horn was opened along the longitudinal line and washed in Ca^2+^- and Mg^2+^-free PBS. The mucosal layer was stripped off, minced, and digested overnight with 0.2% collagenase. Endometrial glands were isolated from the digested tissues by filtration (40 µm pore size) followed by a gravitational sedimentation. The sedimented glands were collected and cultured in DMEM containing 5% FBS, 100 U/mL penicillin, 100 µg/mL streptomycin, 100 µg/mL kanamycin, 1% non-essential amino acids, and 10 µg/mL insulin) at 37°C in 5% CO_2_. The culture media were refreshed every 2 days. The contamination of immune cells was removed, and the attached epithelial cells became confluent monolayers within 3-4 days. The remaining stromal cells were removed by adding 0.02 % collagenase in serum-free medium for 24 h. After reaching 90% confluence, the cells were trypsinized and sub-cultured to an appropriate cell culture vessel for each experiment.

Following our previous study [36], PGE purity over 98% was determined by the immunocytochemistry staining of anti-pan cytokeratin antibody and transepithelial electrical resistance (TER) of greater than 400 Ω.cm^2^. PGE monolayers with TER of 400-800 Ω.cm^2^ which were considered as high tight junction integrity for studying ion transport were chosen for inoculation. The contamination of PGE with *Mycoplasma spp*., swine fever or PRRSV was assessed by a multiplex reverse transcription quantitative polymerase chain reaction (RT-qPCR) detection kit (Microplasma 16 s Ribosomal RNA Gene Genesig® Standard kit, [Primerdesign, Camberley, UK]; Virotype® CSFV RT-PCR kit, [QiagenIAGEN, Leipzig, Germany]) [12]. The mentioned pathogen-negative PGE was chosen for PRRSV inoculation.

### Preparation of PRRSV inoculum

PRRSV was isolated from the lungs of pigs with respiratory and reproductive illness and positive PRRSV sera at the Farm Animal Hospital (Faculty of Veterinary Science, Chulalongkorn University, Nakorn Pathom, Thailand). To confirm and prepare PRRSV inoculum, 2.3 g of the infected lung tissues was minced, homogenized in 15 mL of cold FBS-free DMEM, and centrifuged at 10,000 × g and 4°C for 10 min following the previous protocol [37]. The supernatants were collected and filtered through a 0.2-µm syringe, diluted with FBS-free DMEM at a 1:1 ratio, and freshly proceeded to RT-qPCR using primers specific to ORF7 of type 1/type 2, ORF 7 of type 1, and ORF 7 of type 2, as previously described [12, 38].

According to our previous study [12], 1 µg of cDNA template was mixed with qPCR SYBR master mix in the presence of forward and reverse primers. All reactions were subjected to CFX96™ Real-Time PCR Detection System (Bio-rad, Hercules, CA, USA) using the following cycle: 95°C for 3 min to activate the reaction, followed by 40 cycles of amplification steps, including denaturation at 95°C for 20 s, annealing at 60℃ for 30 s and extension at 72℃ for 30 s, respectively. The specificity of amplified products was confirmed using 1.5% agarose gel electrophoresis and melting curve analysis. The lungs of PRRSV-negative pigs were isolated and used for mock infection. No amplicons were produced in the mock control.

### PRRSV inoculum quantification

All PRRSV inoculum were quantitated using MARC-145 cells (ATCC American Type Culture Collection, VA, USA) cultivating in a 25-cm^2^ flask (Costar^®^, Corning, MA, USA) supplemented with DMEM containing 10% FBS. To quantitate the virus concentration as described previously [12], PRRSV inoculum at 1 mL of 10-fold serial dilutions (10^−1^ to 10^−6^) were incubated with the confluent MARC-145 cells at 37°C in 5% CO_2_ for 1 h. After viral inoculation, the cells were washed and replaced with the fresh media. CPEs in each MARC-145 cell culture well were observed microscopically at 2-, 4- and 6-days post-infection (dpi). Following the Reed-Muench method [39], the dilution that produced CPEs by 50% was considered the tissue culture infective dose 50% (TCID_50_)/mL PRRSV inoculum stock at the endpoint dilution of 10^5^ TCID_50_/mL to produce the pathological changes was used in this study [12].

### PRRSV inoculation

PGE (1×10^6^ cells/mL) were seeded into 24 mm membrane cell culture inserts (Transwell, MA, USA) or 25 cm^2^ flasks (Costar, MA, USA), and maintained in the culture medium for 7 days to become confluence. PGE were then allocated to Mock-, PRRSV type-1 or type-2 group (n = 5 pigs each group). According to our previous protocol [12], the PGE monolayer in 24 mm culture inserts or 25 cm^2^ flasks was respectively inoculated with 1 mL or 5 mL of PRRSV 10^5^ TCID_50_/mL inoculum in 5% CO_2_ at 37°C for 1 h. Inoculation with solution extracted from PRRSV-negative lungs was used for the mock group. After 1 h of viral adsorption, the cells were washed and replaced with fresh medium for 2-6 days. Each inoculation was performed in duplicate.

### Immunohistochemistry, detection, and quantitation of PRRSV infection and mediators

All PRRSV-infected PGE were confirmed by the presence of CPE at 2, 4 and 6 dpi. All CPEs were observed and measured in filter-grown PGE under light microscope with digital camera (BX50F and UC50, Olympus, Tokyo, Japan). The total area of CPE was normalized with the membrane filter area and reported as a percentage. To detect and quantitate PRRSV existent, PRRSV isolated from PGE and culture medium were assessed by RT-qPCR with the inoculation to MARC-145 cells following the Reed-Muench method [39].

Following the detection of CPE in PRRSV-infected PGE, immunostaining with primary antibodies (Ab) to recognize viral protein PRRSV-GP5, and PRRSV mediators CD151, CD163, sialoadhesin (Sn), integrin, and vimentin was performed. Following our previous protocol [12], filter-grown PGE were fixed with 4% paraformaldehyde in PBS, pH 7.4 for 10 min at 25°C. Fixed PGEs were treated with the non-specific blocking solution, 10% H2O2 in methanol, followed by 4% goat serum in PBS for 4 h. The treated samples were incubated with primary antibodies at 4°C overnight followed by universal HRP-conjugated secondary antibodies (Vectastain^®^ Elite ABC-HRP kit, Vector Laboratories, Inc., Burlingame, CA, USA). Immunoreactivity was developed by incubating membrane filter with DAB (3,3-diaminobenzidine tetrahydrochloride) substrate (Sigma Aldrich, MO, USA) and counter-stained with hematoxylin (Invitrogen, Waltham, MA, USA). Immunoreactive area was visualized under light microscope connected to a digital camera (BX50F and UC50, Olympus, Tokyo, Japan) with 20x magnification. All positive cells were captured by the digital images at the magnification of 20x and analyzed by ImageJ software (National Institutes of Health, Bethesda, MD, USA). The area in pixel numbers of immunoreactive cells (dark-brownish staining) was measured and calculated as the percentage of the total area of PGE, which comprised immunoreactive and non-immunoreactive cells per fields. The means of all percentages of positive cells in each group were compared.

### Total RNA isolation and reverse transcription

Total RNA was collected from PGE (uninfected, mock, type 1, and type 2 groups) grown in T25 flask using TRIzol® reagent (Invitrogen, Waltham, MA, USA). The concentration of total RNA was measured for optical density at 260 nm (OD260) using Nanodrop (NanoDrop 2000, Thermo Fisher Scientific, Waltham, MA USA). The purity of RNA was acceptable if the OD260/OD280 ratio was at 1.8-2.0. Reverse transcription was done by using iScript® Select cDNA synthesis Kit (Bio-Rad Laboratories, Hercules, CA, USA). Briefly, a 20 µl of reaction mixture consisting of total RNA 3 µg, oligo dT primer and iScript reaction was prepared. First-strand DNA was synthesized in thermocycler (Biometra GmbH, Göttingen, Germany) using the following cycle: 25°C for 3 min, 46°C for 20 min, and 95°C for 1 min. cDNA concentration was measured using NanoDrop and product stored at −20°C until performing real-time PCR.

### Real-time PCR

The mRNA expression of PRRSV mediators, TLRs, and cytokines was carried out using a GeneOn SYBR green based qPCR kit (GeneOn, Deutschland, Germany) following the previous protocol [12, 20]. Briefly, a 20 µl of PCR reaction containing 1 µg of cDNA template, 2x qPCR SYBR master mix, and a pair of forward and reverse primers for each gene (Table 1) was prepared. Forty cycles of reaction at 95°C, 60°C, and 72°C for 20, 30, and 30 s, respectively, were carried out in a DNA Thermal Cycler (CFX96™, Bio-rad, Hercules, CA, USA). The amplicons were evaluated for the specificity of product by running 1.5% agarose gel electrophoresis and analyzing melting curve. The relative mRNA expression of interested genes was determined by normalization with GAPDH mRNA expression and reported as fold change compared to those of uninfected PGE using 2*^−ΔΔCt^* calculation [40].

### Semiquantitative Western blot analysis

PGE proteins were harvested using lysis buffer containing 50 mM tris, 150 mM NaCl, 1mM EGTA, 1mM PMSF, 1% NP-40, 6.02 mM sodium deoxycholate, 0.01 mg/ml aprotinin, 1 mM NaF and a cocktail protease inhibitor. Cell lysate was centrifuged at 12000 rpm for 15 min at 4°C. The supernatant was collected and measured for a protein concentration using the BCA™ protein assay (Thermo Fisher Scientific, MA, USA). The protein sample was diluted with Laemmli buffer containing β-mercaptoethanol (Bio-rad, Hercules, CA, USA) and incubated at 65°C for 5 min. The obtained denatured protein (30 μg) was separated by 10% sodium dodecyl sulphate-polyacrylamide gel electrophoresis (SDS-PAGE) and blotted to a PVDF membrane (Millipore®, St Louis, MO, USA). After blocking non-specific protein with 2% bovine serum albumin, the blotted membrane was probed with primary antibodies at 4°C for 12 h and further stained with HRP-conjugated secondary antibodies for 1 h at room temperature. Dilution of all antibodies was used following the manufacturer’s instruction. An ECL substrate (Santacruz Biotechnology, Dallas, TX, USA) was used to develop immunoreactive bands, which were visualized by exposure to an X-ray film (GE healthcare, Bloomington, IL, USA). The protein expression of TLRs (n = 5 pigs each group) was analyzed by Scion image software and normalized to β-actin. The results were represented as relative fold changes of protein expression over those of mock group.

### Measurement of cytokine secretion

Media collected from the apical and basolateral compartment of filter-grown PGE at 0, 2, 4 and 6 dpi were determined for cytokine secretion, which consisted of chemokine (C-C motif) ligand 2 (CCL-2), IL-1β, IL-6, IL-8, IL-10, TNF-α, IFN-α and IFN-γ by enzyme-linked immunosorbent assay (ELISA) using commercial kits (R&D System, Minneapolis, MN, USA). The ELISA processes were performed according to the instruction of commercial kits (n = 5 pigs for each group). Cytokine concentration was measured at OD450, by a microplate reader (Biotek, Winooski, VT, USA), in which a background OD620 was subtracted from all data. The concentration was calculated from a standard curve. The accumulated concentrations of each cytokine from 0-6 dpi were normalized by dividing with those of uninfected cells and displayed as fold changes of uninfected cells.

### Statistical analysis

All PGE data were obtained from 5 pigs per group and shown as the mean ± SEM. Statistical analysis was done by GraphPad Prism 9.0 (GraphPad software Inc., San Diego, CA, USA). The percentages of CPE and PRRSV positive cells among different dpi were analyzed by two-way ANOVA, and the mRNA expression of interested genes, protein expression, and cytokine secretion were analyzed by one-way ANOVA. The multiple comparison with the uninfected group was performed by Bonferroni’s post hoc test. A *p* value of less than 0.05 was considered as significant differences.

## Acknowledgments

The authors thank the government qualified slaughterhouse in Bangkok, Thailand. for providing samples used in this experiment. Department of Physiology and the Department of Medicine, Faculty of Veterinary Science, Chulalongkorn University and the Department of Physiology, Faculty of Medicine, Srinakharinwirot University for facilities used in this study.

